# The role of muscle fascia in heterotopic ossification and maintenance of skeletal muscle integrity in fibrodysplasia ossificans progressiva

**DOI:** 10.1101/2025.11.12.688008

**Authors:** L. Russell Hanson, Katherine L. Scalise, Rayna M. Esch, Liv Nevo, David J. Goldhamer

## Abstract

The rare genetic disorder fibrodysplasia ossificans progressiva (FOP) is characterized by progressive heterotopic ossification (HO) of skeletal muscles and associated soft tissues. FOP is caused by a gain-of-function mutation in the type l BMP receptor ACVR1 (ALK2) that renders the receptor inappropriately responsive to activin ligands. HO is associated with muscle destruction and compromised muscle regeneration, although little is known of the mechanistic relationship between these pathophysiological disease manifestations. In mouse FOP models, HO is experimentally induced by direct injury to muscle using chemical or mechanical means, thereby obscuring the relationship between HO formation and muscle destruction. We show that direct muscle injury is not required for induction of a robust HO response. Rather, a small incision in the fascia superior to the tibialis anterior muscle was sufficient to induce HO when fibro-adipogenic progenitors (FAPs) were targeted for *Acvr1^R^*^206^*^H^*expression. Intermuscular fascial layers were the primary sites of lesional growth when HO was exacerbated by genetic, pharmacological, or physical means. In contrast to control mice, fascial injury in FOP mice caused pronounced destruction of the muscle subjacent to the injured fascia. Further, areas of muscle degeneration did not undergo a productive regenerative response. Unlike most models of impaired regeneration, adipocyte accumulations were not observed in areas of muscle degeneration, which were destined for pathological bone formation. These data point to the primary role of fascia in HO initiation and growth, and indicate that *Acvr1^R^*^206^*^H^*-expressing FAPs directly or indirectly create an abnormal tissue environment that destabilizes muscle tissue and is incompatible with muscle regeneration. The advantages of this new model of injury-induced HO for understanding early events in FOP pathogenesis are discussed.

**One sentence summary:** Muscle fascia serves a primary role in FOP pathogenesis.

## Introduction

Fibrodysplasia ossificans progressiva (FOP) is a rare autosomal dominant musculoskeletal disease characterized by heterotopic ossification (HO) of skeletal muscle and connective tissues, typically beginning in early childhood. The cumulative effects of HO result in muscle loss, ankylosis of major joints, and other complications that cause pain, severely restrict mobility, and reduce life expectancy^1–3^. The great majority of cases of FOP with classical clinical features result from an arginine to histidine (R206H) substitution in the highly conserved glycine-serine domain of the bone morphogenetic protein (BMP) type I receptor, ACVR1 (also known as ALK2)^4–6^. This mutation renders ACVR1 responsive to the TGFβ ligand activin A^7–10^, inappropriately activating osteogenic signaling in tissue-resident mesenchymal progenitors that become co-opted for skeletogenesis. Soft tissue trauma can induce HO formation, and even mild injury (e.g., immunizations, incidental bumps and bruises) can cause severe HO^3,11^. In addition, individuals with FOP often experience spontaneous bouts of HO that are not associated with a known proximate trigger^7,8^.

Muscle and connective tissue loss are well-described clinical features of FOP pathophysiology^11–13^, although the mechanistic underpinnings of this tissue destruction are poorly understood. Skeletal muscle is a highly regenerative tissue owing to the action of muscle stem cells (satellite cells), which are activated from quiescence by injury stimuli. Despite muscle damage, few regenerated muscle fibers were observed in areas of HO in a genetically accurate FOP mouse model^14^, indicating that a diminished regenerative response may contribute to muscle loss. In this model, expression of *Acvr1^R^*^206^*^H^*was targeted to fibro-adipogenic progenitors (FAPs), a known cell-of-origin of HO in FOP models^14–18^. Under normal physiological conditions, FAPs support regeneration by promoting satellite cell proliferation and differentiation^19–23^, and *Acvr1^R^*^206^*^H^* might function cell non-autonomously in FAPs to impair satellite cell functions in FOP. Alternatively, or in addition, evidence from cell culture experiments indicates that *Acvr1^R^*^206^*^H^* affects satellite cell function cell-autonomously^24^.

The fascial system is a body-wide network of interconnected soft connective tissues that serve many functions in maintaining tissue homeostasis, including responses to stretch and mechanical load, immunological responses to infection, wound healing, and others^25,26^. Deep fascia of the limbs is essentially dense connective tissue that separates muscles into compartments (e.g., anterior vs. posterior in the lower limb) and aids in transmitting force between muscles within a compartment to coordinate muscle activity. Additionally, each muscle and its component muscle fibers are surrounded by fascia containing loose connective tissue: the epimysium and endomysium, respectively. Injury-induced HO in FOP mouse models is generally assumed to originate intramuscularly, in contrast to spontaneous HO, which primarily affects tendons, ligaments, and joints, and tends to follow fascial planes^14,16,27^. In injury models, muscle injury is typically induced by intramuscular injection of a myotoxin or by physical trauma to muscle tissue^14,15,17,27^. However, our previous work showed that muscle injury usually results in HO within intermuscular fascia rather than originating intramuscularly^15,17^. In addition, a “sub-threshold” injury modality was identified whereby intramuscular injection of methylcellulose elicited a localized muscle regenerative response but did not induce FAP-driven HO, despite the widespread distribution of FAPs within the muscle endomysium^15^. These latter data suggest that muscle injury per se is not a sufficient stimulus to induce HO.

To assess the role of fascial injury in HO induction and to define the effects on muscle integrity and regeneration, we used genetic mouse models of FOP in which *Acvr1^R^*^206^*^H^*was conditionally expressed in FAPs using Tie2-Cre^28^ or *Pdgfra^CreERT^*^2^ ^29^ drivers. We show that a small incision to the epimysium and superficial fascia that avoids direct injury to the underlying muscle is sufficient to drive HO with high penetrance. Unlike other injury modalities, HO resulting from fascial injury developed in a predictable anatomical location, which was superficial to, but encroaching on, the underlying muscle. Fascial injury also causes directional and progressive muscle destruction, beginning in proximity to the overlying fascial incision, and the muscle fails to regenerate. Surprisingly, unlike other models of impaired regeneration^30–33^, failed regeneration in this model is not accompanied by accumulations of adipose tissue. These results demonstrate a role for fascia-resident FAPs in HO development and muscle loss and provide a model to interrogate FAP-satellite cell interactions in FOP.

## Materials and Methods

### Mouse Models, Breeding, and Genotyping

Standard breeding schemes were used to generate experimental animals. Control mice were generated from the same crosses and contained only the Cre allele. Cre drivers were introduced through the male parental germline to avoid global recombination, which can occur when Cre is introduced through the female germline^34,35^. Except where noted, female experimental mice were used for all studies because HO penetrance ^34,36,37^ and average volumes are greater than in males, as shown previously^18^ and documented here. The following knockin and transgenic mouse lines were used: *Acvr1^tnR^*^206^*^H^* ^14^, the Cre drivers Tie2-Cre (JAX #008863)^28^, *Pdgfra^CreERT^*^2^ (JAX #032770)^29^, and *Pax7^CreERT^*^2^ (JAX #017763)^38^, the *Pdgfra*-driven GFP reporter *Pdgfra^H2B-EGFP^* (JAX#007669)^39^, a Cre-dependent GFP reporter *R26^NG^*(JAX#012429)^40^, conditional knockout alleles *Acvr1^flox^* ^41^, *Myf5^loxP^* ^42^, and *MyoD^L2G^* ^43^, and the constitutive knockout allele *MyoD^Neo^*(*MyoD^m^*^1^)^44^. Experimental mice used in this study were 8-12 weeks old and maintained on an enriched FVB background. Genotyping by PCR was conducted as described previously^17,38,42,43^.

### Tamoxifen, Cardiotoxin, and Antibody Injections

Tamoxifen (Toronto Research Chemicals, T006000) was injected intraperitoneally (IP) at 200 mg/kg from a 20 mg/ml working solution. For dosing of *Acvr1^tnR^*^206^*^H/+^*;*Pdgfra^CreERT^*^2^*^/+^* mice, tamoxifen was prepared in sunflower seed oil (Sigma-Aldrich, 8001-21-6), and mice were injected for three consecutive days. For *Pax7^CreERT^*^2^*^/+^*;*MyoD^L2G/neo^*;*Myf5^loxP/loxP^*;*R26^NG/+^*mice, tamoxifen was prepared in corn oil (Sigma-Aldrich, C8267), and mice received five injections on an every-other-day schedule over 9 days. For both mouse strains, fascial injury was performed after a 5-day washout period.

To induce direct muscle injury, *Naja Pallida* cardiotoxin (Latoxan, L8102) was injected into the tibialis anterior (TA) muscle as previously described^17^. Muscle regeneration was assessed at 6 days post-injury (dpi).

To pharmacologically exacerbate HO, *Acvr1^tnR^*^206^*^H/+^*;Tie2-Cre mice were injected IP with the anti-ACVR1 antibody, JAB0505, at a dose of 10 mg/kg immediately prior to surgery, as previously described^45^.

### µCT Imaging and HO Quantification

HO volume was quantified using the IVIS-Spectrum CT model 128201 (PerkinElmer). µCT images were captured in medium resolution acquisition mode for limb imaging (75 μm voxel size; estimated radiation dose of 132 mGy; 210 s scan time). Mice were imaged under isoflurane anesthesia. The HO volumes were segmented and quantified using 3D Slicer (http://www.slicer.org).

### Fascial Injury

Before surgery, mice were examined and scanned by µCT to verify the absence of HO preceding injury. Experimental and control mice were anesthetized using isoflurane. Limbs were shaved and sterilized using iodine surgical solution (Avrio Health, 67618-150-01), iodine surgical scrub (Avrio Health, 67618-151-17), and 70% ethanol. With the aid of a stereomicroscope, a 5-7 mm incision was made into the skin superficial to the TA muscle. For the standard fascial injury, the fascia was gently lifted at the TA mid-belly using fine forceps, and a 2-3 mm longitudinal incision at the mediolateral midpoint was made in the fascia layers using sharp dissecting scissors. For the long fascial injury, a 7-8 mm incision was made in the fascia along the longitudinal axis at the mediolateral midpoint of the TA. After surgery, the skin was sealed with wound glue (3M-Tissue Adhesive, 70200742529), and mice were singly housed for the duration of the experiment.

### Fixation, Decalcification, Cryopreservation, and Sectioning

Mice were exposed to isoflurane and then cervically dislocated at experimental endpoint. Experimental and control hindlimbs were harvested and fixed in 4% paraformaldehyde (Electron Microscopy Sciences, 19120) for 2 days at 4°C. Limbs were then placed in 12% EDTA (pH 7.2) (Research Products International, E57020) for 2-4 weeks at room temperature and tested periodically for bone decalcification. Limbs were then washed with PBS overnight and placed in 30% sucrose (Sigma-Aldrich, 57-50-1) for 1 day, both at room temperature. Limbs were patted dry, embedded in O.C.T. Compound (Fisher HealthCare, 4585), and frozen in liquid nitrogen-cooled isopentanes (Sigma-Aldrich, M32631). Tissue was stored at -80°C until sectioning. Cryosections were taken at 10-20 μm and mounted on Superfrost Plus slides (Fisher, 1255015), dried overnight at room temperature, and stored at -20°C.

### Histochemistry

For hematoxylin and eosin (H&E) staining, slides were first brought to room temperature and washed in PBS to remove O.C.T. Compound. Slides were then transferred into Mayer’s hematoxylin (Sigma Aldrich, MHS1), then rinsed in deionized water and transferred to 0.1% ammonium hydroxide (Millipore Sigma, 1336-21-6) to blue and mordant the hematoxylin. Slides were progressively dehydrated in graded ethanol solutions and stained with 0.25% eosin Y (Sigma Aldrich, E4382) in 95% ethanol. Slides were then rinsed in 95% ethanol, further dehydrated in 100% ethanol, cleared in xylenes, and mounted with Permount solution (Fisher Chemical, SP15500).

RGB trichrome staining (picrosirius red, fast green, alcian blue) was performed as previously described^46^, with minor modifications. Briefly, slides were washed with PBS for 5 min to dissolve O.C.T. Compound and subsequently fixed in Bouin’s solution for 1 hr at 56°C. After rinsing with tap water, slides were sequentially stained with 1% alcian blue 8GX (pH 2.5) for 20 min, 0.04% fast green FCF for 20 min, and picrosirius red for 30 min, with tap water rinses after each staining step. Finally, slides underwent three 4-min washes in 3% acetic acid before being dehydrated and mounted as above.

### Immunofluorescence

After removing O.C.T. Compound as above, frozen sections were permeabilized with 0.1% Triton-X 100 (Sigma, T8787) in PBS (permeabilization buffer) and blocked in 1% bovine serum albumin (BSA) (Sigma Life Sciences, A2153), 10% goat serum (Sigma Aldrich, G6767), and 0.1% Tween 20 (Sigma, 9005-64-5) in PBS (blocking buffer). Slides were incubated with primary antibody in blocking buffer overnight at 4°C or at room temperature for 2 hr. Rabbit anti-SOX9 (Millipore, AB5535) and rabbit anti-Perilipin (Sigma Aldrich, P1873) antibodies were both used at dilutions of 1:1000. Slides were subsequently washed three times in PBS and incubated with an Alexa-fluor 647 conjugated goat anti-rabbit secondary antibody (Invitrogen, A21245) at room temperature for 1 hr at dilutions of 1:500 (SOX9) or 1:1000 (Perilipin). Slides were then washed three times with PBS, counterstained with DAPI (Roche Diagnostics, 10236276001), washed three times with PBS, and coverslipped using Fluoro-Gel (Electron Microscopy Sciences, 17985-11).

Prior to Dystrophin staining, cryosections were subjected to antigen retrieval as described previously^43^. Subsequently, sections were incubated overnight at 4°C with undiluted mouse anti-Dystrophin antibody supernatant (Developmental Studies Hybridoma Bank, AB_2618170) in 20% FBS (Gibco, S11150) with 1% penicillin-streptomycin in DMEM (Gibco, 11995-065). Slides were washed three times in PBS and incubated with an Alexa-Fluor 647 conjugated goat anti-mouse secondary antibody (Invitrogen, A21236) in blocking buffer at a concentration of 1:1000 at room temperature for 1 hr and processed as described above.

### Microscopy and Imaging

Both brightfield and immunofluorescent images were acquired using the Leica Thunder Imager (Leica DM6 B microscope). The microscope is equipped with four fluorescent filter cubes (excitation wavelengths of 405, 470, 545, and 620 nm) and fluorescent images were captured with a Leica K5 sCMOS monochrome camera. Brightfield images were obtained with a Leica DMC5400 CMOS color camera. All image capture was performed with LAS X software (Leica) with additional image processing performed with both LAS X and FIJI^47^.

To computationally subtract background fluorescence, control slides from mice not incubated with secondary antibody, or lacking GFP or tdTomato, were imaged under the same parameters as experimental slides. Mean grey value was quantified for each channel and the values obtained from control images were subtracted from experimental images using FIJI.

### Myofiber Quantification

The lower hindlimbs of fascial-injured, age-matched *Acvr1^tnR^*^206^*^H/+^*;Tie2-Cre and injured and uninjured control mice were collected at both 6 and 14 dpi (n = 4 per time point for each group). After tissue processing as above, lower hindlimbs were serially cryosectioned transversely and representative sections were stained for Dystrophin and examined for HO. For each mouse, muscle fibers were counted on a section corresponding to the position of maximal HO. As fiber numbers vary substantially along the longitudinal axis of the TA muscle, the comparable anatomical position along the longitudinal axis of the lower hindlimb was used for fiber counts of control mice. Sections were imaged and the images were processed in FIJI to crop out all but TA and extensor digitorum longus (EDL) muscle fibers. Image files were uploaded to Cellpose^48^ and used to train an AI model within Cellpose to automate segmentation and fiber counts. Each image was quality controlled and uncounted myofibers were added manually (< 1% of total fibers). The average myofiber counts of the injured control and experimental mice were normalized to the uninjured controls and represented as a percentage.

### Statistics

Statistical analysis was performed using GraphPad Prism (GraphPad). All values are represented as the mean value +/- the standard error of the mean (SEM). Bone volume comparisons between male and female mice, the standard and long fascial injury, and between *Acvr1^tnR^*^206^*^H/+^*;Tie2-Cre and *Acvr1^tnR^*^206^*^H/+^*;*Pdgfra^CreERT^*^2^*^/+^* mice were performed using an unpaired t-test. Comparison of longitudinal bone growth in long fascial injury mice were performed using a paired t-test. Statistical comparisons of myofiber numbers used a one-way ANOVA with Tukey’s multiple comparisons. Normalization of fiber numbers to uninjured controls (described above) reduces variance in uninjured control group to zero, which violates the ANOVA assumption of homoscedasticity. However, simulations show that when three or more groups are compared using ANOVA followed by post-hoc tests, normalization in this way does not increase the type 1 error rate^49^. Longitudinal studies of bone volumes at 14, 21, and 28 dpi also used ANOVA.

## Results

### Cutting of the superficial fascia and epimysium does not damage the underlying skeletal muscle

After making a small skin incision to expose the TA muscle, the superficial fascia and epimysium at the muscle mid-belly were gently lifted away from the muscle with a fine forceps so that a 2-3 mm cut could be made in the connective tissue layers while avoiding contact with the underlying TA muscle (Supplemental Fig. 1A, B). Histological analysis of control mice (lacking the *Acvr1^tnR^*^206^*^H^*allele) showed no evidence of muscle damage through 14 dpi (Fig. 1A-D). Specifically, the underlying muscle juxtaposed to the site of fascial injury was indistinguishable from uninjured controls, lacking both necrotic fibers and hypercellularity that typify early post-injury stages after direct muscle damage. Further, regenerated fibers were not observed at any post-injury time point (data not shown). In contrast, when approximately 1 mm of the TA muscle of control mice just beneath the site of fascial injury was intentionally cut, fiber loss and hypercellularity were evident by 3 dpi, and regenerated fibers, identified by their central nucleation, were apparent by 6 dpi (Fig. 1E, F). Satellite cell-targeted inactivation of the muscle regulatory genes, *MyoD* and *Myf5*, results in regenerative failure, and the injured muscle exhibits marked hypercellularity and extensive fibrotic and adipogenic accumulations^43^. We performed fascial injury on these double knockout mice under the assumption that even minor injury to the muscle would cause abnormal tissue changes detectable by histology^29^. At 14 dpi, double knockout mice were virtually indistinguishable from controls, with only a slight degree of hypercellularity at the most peripheral aspect of the TA muscle, directly beneath the site of fascial injury. No indications of muscle fiber damage or histological manifestations of abortive regeneration were evident (Fig. 1G, H). Taken together, these data indicate that the fascial injury method employed does not directly damage underlying muscle fibers.

**Figure 1.**
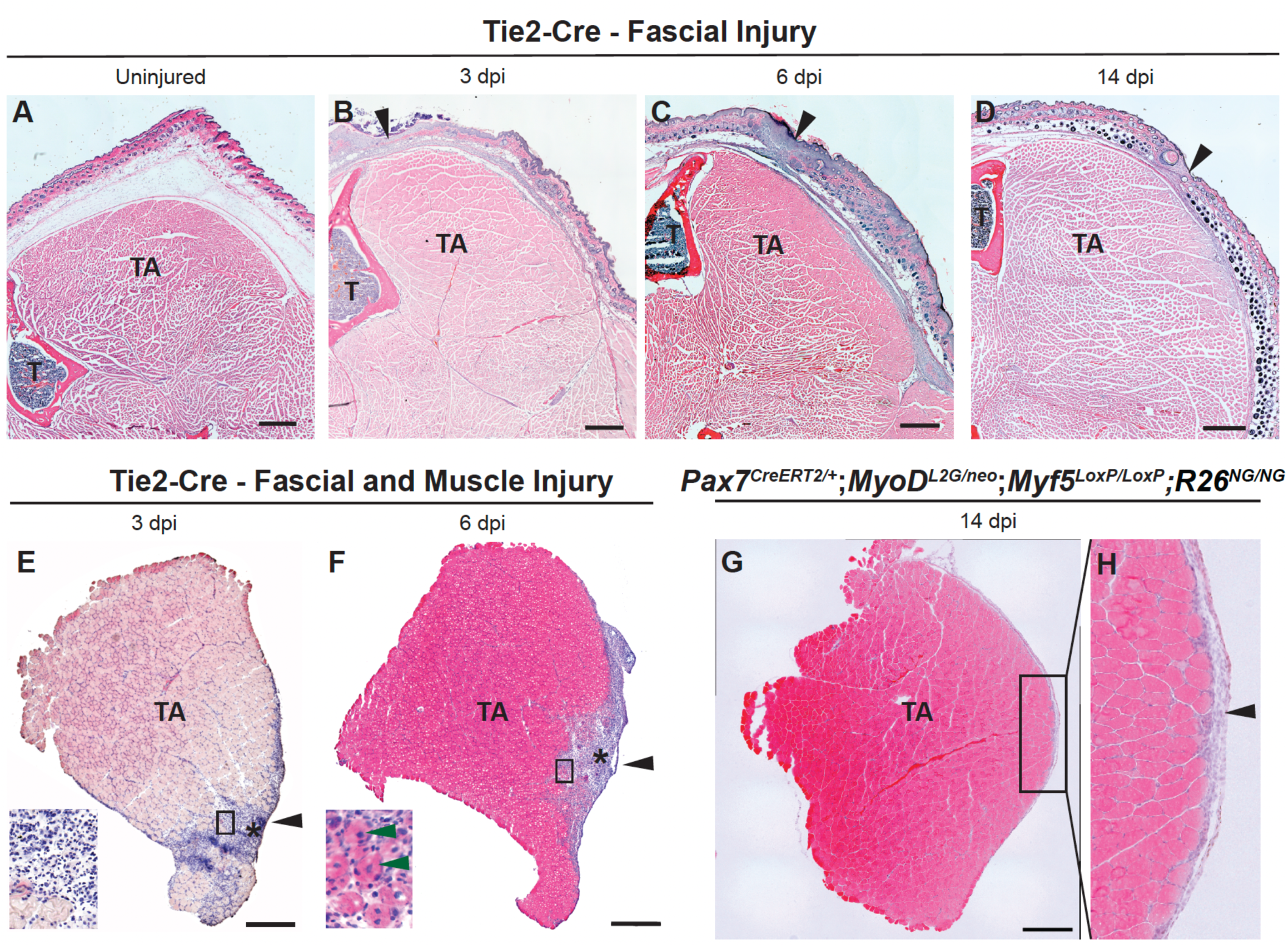
Fascial injury does not damage the underlying skeletal muscle. **(A-D)** Transverse cryosections of the anterior portion of the lower hindlimb of uninjured (A) and fascial-injured (B-D) control mice (lacking *Acvr1^tnR^*^206^*^H^*; n = 4 mice per time point). At all time points post injury, the TA muscle remained intact below the area of fascial injury (arrow) and was histologically similar to uninjured animals (A), with no evidence of muscle damage or fiber regeneration. (**E-F)** Cryosections of TA muscles from fascial-injured control mice in which the TA muscle was purposely incised at a depth of approximately 1 mm (n = 3 mice per time point). At 3 dpi (E), the area of muscle damage (asterisk) was marked by hypercellularity and necrosing myofibers (inset). Nascent muscle fibers (identified by central nucleation) were evident by 6 dpi (F, green arrowheads in inset). **(G, H)** Cryosection of the TA muscle from a *Pax7^CreERT^*^2^*^/+^*;*MyoD^L2G/Neo^*;*Myf5^loxP/loxP^*;*R26^NG/+^* mouse at 14 dpi (n = 2 mice). Panel (H) represents the boxed area in (G). Mice showed a slight increase in cellularity at the site of the fascial incision, but with no indication of muscle damage. Black arrowheads in (B, C, D, E, F, H) mark the approximate site of fascial injury. T, tibia; TA, tibialis anterior. All sections were stained with H&E. Scale bars in (A-G) = 500 µm.

### Fascial injury is sufficient to induce HO in FOP mice

We previously showed that targeting FAPs for *Acvr1^R^*^206^*^H^* expression using the Tie2-Cre driver^28^ induces robust HO following muscle injury elicited by cardiotoxin injection or pinch injury^9,17,45^. Notably, spontaneous HO in *Acvr1^tnR^*^206^*^H/+^*;Tie2-Cre mice tends to follow fascial planes^9^. Additionally, intramuscular injection of methylcellulose into *Acvr1^tnR^*^206^*^H/+^*;Tie2-Cre mice caused a localized regenerative response but was not sufficient to induce HO^9,45^, indicating that muscle injury per se is not sufficient to induce HO. These observations prompted us to examine whether injury to the fascia, without direct injury to skeletal muscle, could trigger an HO response in *Acvr1^tnR^*^206^*^H/+^*;Tie2-Cre mice. The fascia of both male and female *Acvr1^tnR^*^206^*^H/+^*;Tie2-Cre mice was cut as described above, and mineralized HO volume quantified by μCT at 14 dpi, as previously described^14^. HO was induced in all but two mice, suggesting that direct muscle injury is not necessary to initiate HO formation (Fig. 2A, C; Supplemental Fig. 2A; n = 23). Skin incision alone (sham-operated control) can result in HO, but at lower penetrance (Supplemental Fig. 1C, D); HO caused by skin incision is clearly distinguishable from fascial injury-induced HO by its anatomical position and lack of effect on the underlying musculature (Supplemental Fig. 1D, and see below). Of note, after fascial injury, females exhibited a greater average HO volume and higher variability in HO response than males (Fig. 2A), as previously reported using both cardiotoxin- and pinch-induced muscle injury as stimuli^18^. We confirmed FAP-mediated HO induction using a *Pdgfra^CreERT^*^2^ driver ^29^ (Fig. 2B, D), which targets FAPs with greater specificity than Tie2-Cre^21,50–52^.

**Figure 2.**
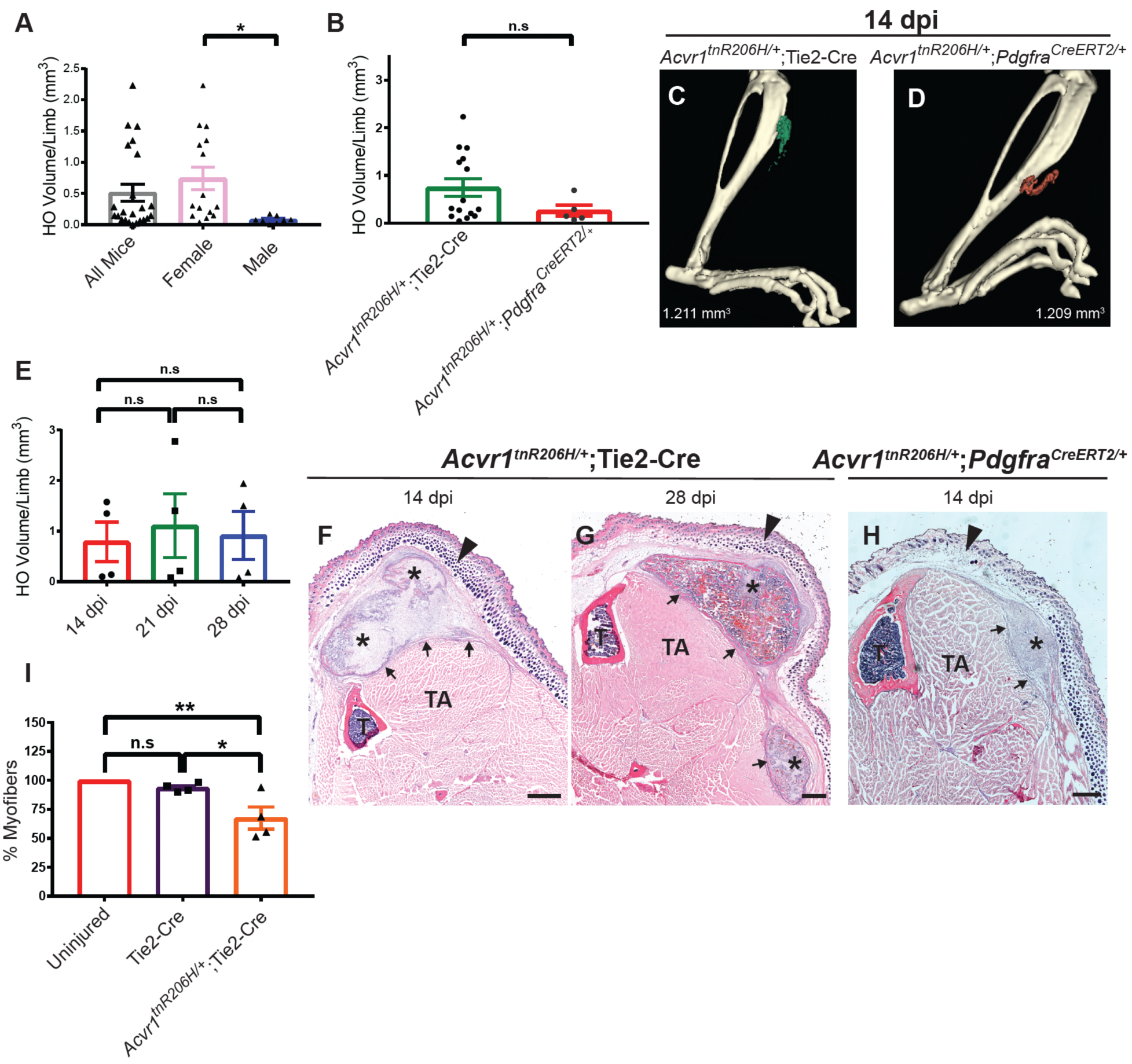
Fascial injury drives HO formation and muscle fiber loss in FOP mice. **(A)** Quantification of HO volume in *Acvr1^tnR^*^206^*^H/+^*;Tie2-Cre mice at 14 dpi (n = 23). Male mice (n = 8) displayed a lower average bone volume after fascial injury than female mice (n = 15). **(B)** Comparison of bone volumes between *Acvr1^tnR^*^206^*^H/+^*;Tie2-Cre (n = 15) and *Acvr1^tnR^*^206^*^H/+^*;*Pdgfra^CreERT^*^2^*^/+^* (n = 5) mice. **(C, D)** Representative µCT images of HO in *Acvr1^tnR^*^206^*^H/+^*;Tie2-Cre (pseudocolored green) and *Acvr1^tnR^*^206^*^H/+^*;*Pdgfra^CreERT^*^2^*^/+^* (pseudocolored maroon) mice at 14 dpi. **(E)** Longitudinal analysis of HO growth in *Acvr1^tnR^*^206^*^H/+^*;Tie2-Cre mice through 4 weeks post-injury. **(F-H)** H&E-stained cryosections of the anterior portion of the lower hindlimb from fascial-injured *Acvr1^tnR^*^206^*^H/+^*;Tie2-Cre (F, G) and *Acvr1^tnR^*^206^*^H/+^*;*Pdgfra^CreERT^*^2^*^/+^* (H) mice. HO (asterisks) developed near the site of fascial injury with HO occupying space formerly occupied by myofibers. The boundary between muscle tissue and HO lesions is demarcated with arrows. Black arrowheads in (F-H) mark the approximate site of fascial injury. **(I)** Myofiber quantification at 14 dpi. Control and *Acvr1^tnR^*^206^*^H/+^*;Tie2-Cre FOP mice were compared to uninjured FOP mice. n = 4 for each condition. T, tibia; TA, tibialis anterior. Except for (A), only female mice were used. Scale bars in (F-H) = 500 µm. Error bars in (A, B, E, I) represent ± SEM. *p ≤ 0.05; **p ≤ 0.01.

To define the temporal trajectory of skeletogenesis in this new HO induction model, we conducted a longitudinal study of HO formation in *Acvr1^tnR^*^206^*^H/+^*;Tie2-Cre FOP mice using μCT imaging at days 14, 21, and 28 after fascial injury (Fig. 2E). This and subsequent analyses used only females, as the more robust and consistent response facilitated phenotypic characterization and increased the statistical power of the analysis. Following muscle pinch injury or cardiotoxin injection, average mineralized bone volume reaches a maximum at approximately 14 dpi, although longitudinal analysis of individual mice often showed modest growth or bone resorption through 28 dpi^15,18^. The kinetics of HO formation following fascial injury were similar to these other models, with most mineralized bone formation occurring by 14 dpi (Fig. 2E). As with other injury modalities, HO volumes between female mice varied substantially (Fig. 2A, E; Supplemental Fig. 2B).

Histological analysis was employed to define the anatomical location of skeletal lesions in relation to the site of fascial injury and the underlying TA muscle. Following μCT imaging at 14 and 28 dpi, whole lower hindlimbs were fixed, decalcified, cryosectioned, and stained with hematoxylin and eosin (H&E). Whereas the anatomical location of HO lesions following cardiotoxin injection or muscle pinch injury can be highly variable, fascial injury consistently induced HO in a defined and reproducible location in close proximity to the injury site (Fig. 2F-H). Interestingly, unlike sham-operated controls (Supplemental Fig. 1D), HO lesions impinged on the TA muscle such that a portion of the anatomical compartment normally occupied by TA muscle fibers contained lesional tissue (Fig. 2F-H). TA muscle fiber numbers were quantified to distinguish whether the impingement on the TA muscle represented its compression by lesional tissue or the loss of TA muscle fibers. At 14 dpi, there was a significant reduction in the number of TA muscle fibers in FOP mice compared to injured and uninjured control mice (Fig. 2I). Collectively, these data demonstrate that HO can be induced in the absence of direct muscle damage, and that a small cut to the superficial fascia and epimysium is sufficient to induce a replicable HO response at a predictable anatomical location. Further, expression of *Acvr1^R^*^206^*^H^* in FAPs directly or indirectly leads to muscle fiber loss.

### Fascial injury causes degeneration of subjacent musculature in FOP mice

Although fascial injury had no obvious effect on the underlying muscle in control mice (Fig. 1), the loss of muscle tissue in *Acvr1^tnR^*^206^*^H/+^*;Tie2-cre mice at endpoint led us to investigate whether fascial injury and the consequent development of HO impacts muscle integrity in FOP mice. The lower hindlimbs of control and FOP mice were collected at 0, 1, 3, 6, and 10 days after fascial injury, and histological sections were prepared as above. At 0 (Fig. 3A) and 1 dpi (not shown), the TA muscle of FOP mice was indistinguishable from injured and uninjured controls (Fig. 1A-C), with no evidence of fiber necrosis or cellular infiltrates that are characteristic of cardiotoxin-injured muscle at early time points^53^. At 3 dpi, however, FOP mice exhibited apparent fiber loss and hypercellularity at the antero-lateral aspect of the TA muscle, in close proximity to the fascial injury (Fig. 3B). Hypercellularity and apparent muscle fiber loss were more extreme at 6 and 10 dpi (Fig. 3C, D). At these stages, the area of muscle fiber loss and hypercellularity comprised approximately one-third of the normal TA muscle compartment (Fig. 3C-E). Interestingly, muscle degeneration was self-limiting, with the area of muscle loss forming a strict boundary with apparently unaffected areas of the TA muscle (Fig. 3C, D; Fig. 4B, C). For some analyses, the *Pdgfra^H2B-eGFP^*reporter^54^ was included to label FAPs in proximity to the site of injury. In uninjured animals, FAPs were abundant in the fascia superficial to the TA muscle and in the skin, and were interspersed in the TA muscle endomysium (Fig. 3F). By 6 dpi, the hypercellular area contained a substantial number of GFP+ cells (presumptive FAPs; approximately 40% of mononuclear cells), and many GFP+ cells expressed the SRY-Box Transcription Factor 9 (SOX9) (Fig. 3G, H), an essential regulator of cartilage determination and differentiation^55^. By 10 dpi, most SOX9+ cells were no longer GFP+ (Fig. I, J), likely reflecting the downregulation of *Pdgfra* as skeletal lesions matured.

**Figure 3.**
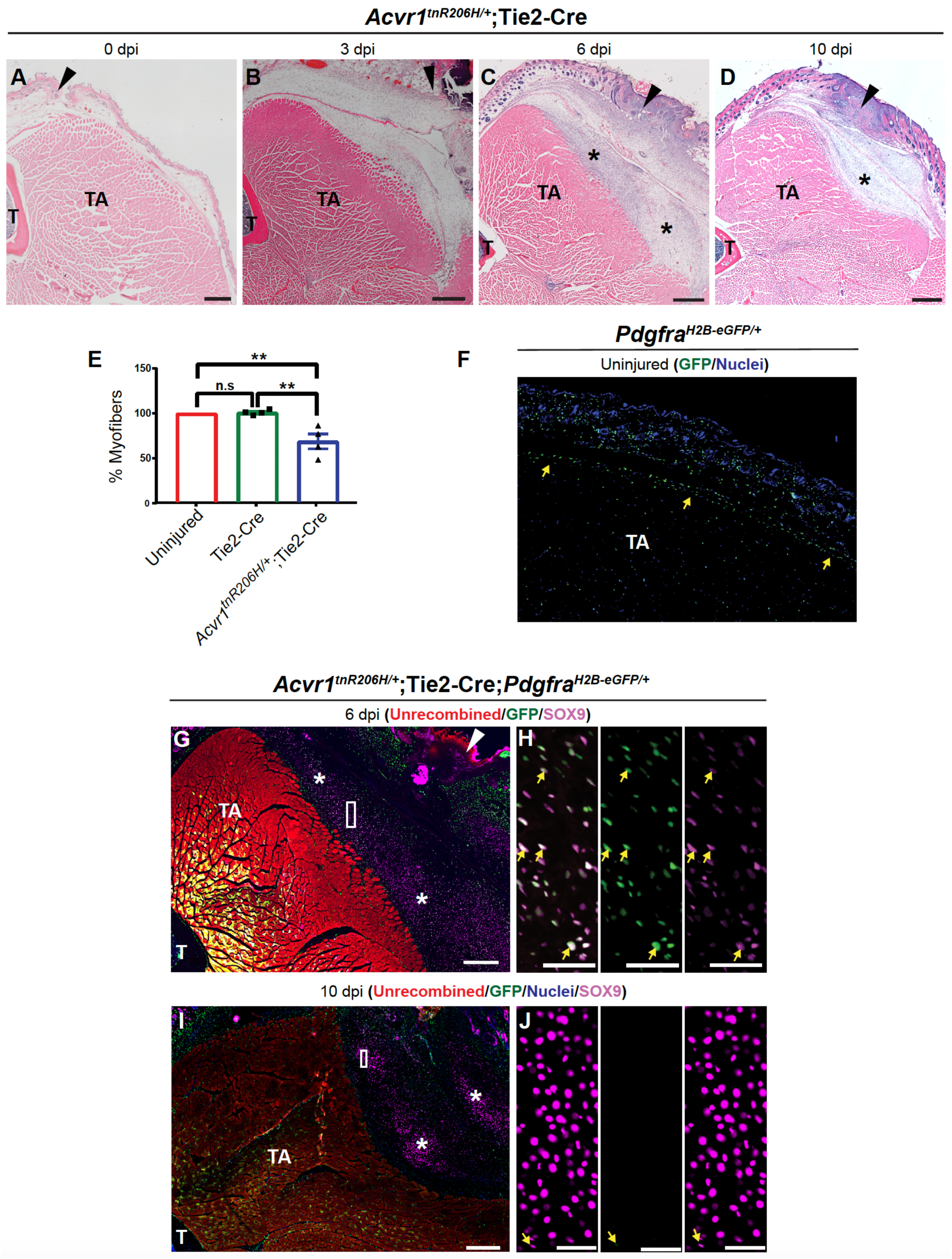
Muscle loss is evident at pre-skeletal lesional stages. **(A-D)** H&E-stained cryosections of the anterior portion of fascial-injured *Acvr1^tnR^*^206^*^H/+^*;Tie2-Cre mice at 0, 3, 6, and 10 dpi (n = 4 for each time point). Like control animals (Fig. 1), FOP mice did not display signs of muscle damage directly after injury. By 3 dpi (B), FOP mice showed increased hypercellularity below the site of injury with an apparent loss of TA and EDL muscle fibers. Muscle destruction and hypercellularity had increased by 6 (C) and 10 (D) dpi. **(E)** Quantification of muscle fiber loss in the TA and EDL muscles at 6 dpi (n = 4). Error bars represent ± SEM. **p ≤ 0.01. **(F)** Fluorescent image of a cryostat section from an uninjured *Pdgfra^H2B-EGFP/+^* mouse. GFP+ FAPs were most abundant in the epimysium/superficial fascia (arrows) and skin. FAPs were also interspersed throughout the TA interstitium. **(G, H)** Fluorescent images of a cryosection from an *Acvr1^tnR^*^206^*^H/+^*;Tie2-Cre;*Pdgfra^H2B-EGFP/+^* mouse at 6 dpi. The panels in (H) correspond to the boxed area in (G). GFP+ cells were abundant in hypercellular regions formerly occupied by muscle fibers, and many GFP+ cells stained positively for SOX9 (H; examples are shown at arrows). The left panel in (H) is a merged image showing colocalization of GFP and SOX9. The middle and right panels are single-channel images. **(I, J)** Fluorescent images of a cryosection from an *Acvr1^tnR^*^206^*^H/+^*;Tie2-Cre;*Pdgfra^H2B-EGFP/+^* mouse at 10 dpi. The panels in (J) correspond to the boxed area in (I). As pre-skeletal lesional tissue matured, fewer SOX9+ cells expressed GFP. Only one cell weakly positive for GFP is identifiable in the boxed area (arrow, middle panel of J). Arrowheads in (A-D, G) mark the approximate site of fascial injury. Asterisks in (C, D, G, I) denote areas of hypercellularity. Only female mice were used. T, tibia; TA, tibialis anterior. Scale bars in (A-C, F, H, I) = 500 µm, Scale bars in (G, J) = 50 µm.

**Figure 4.**
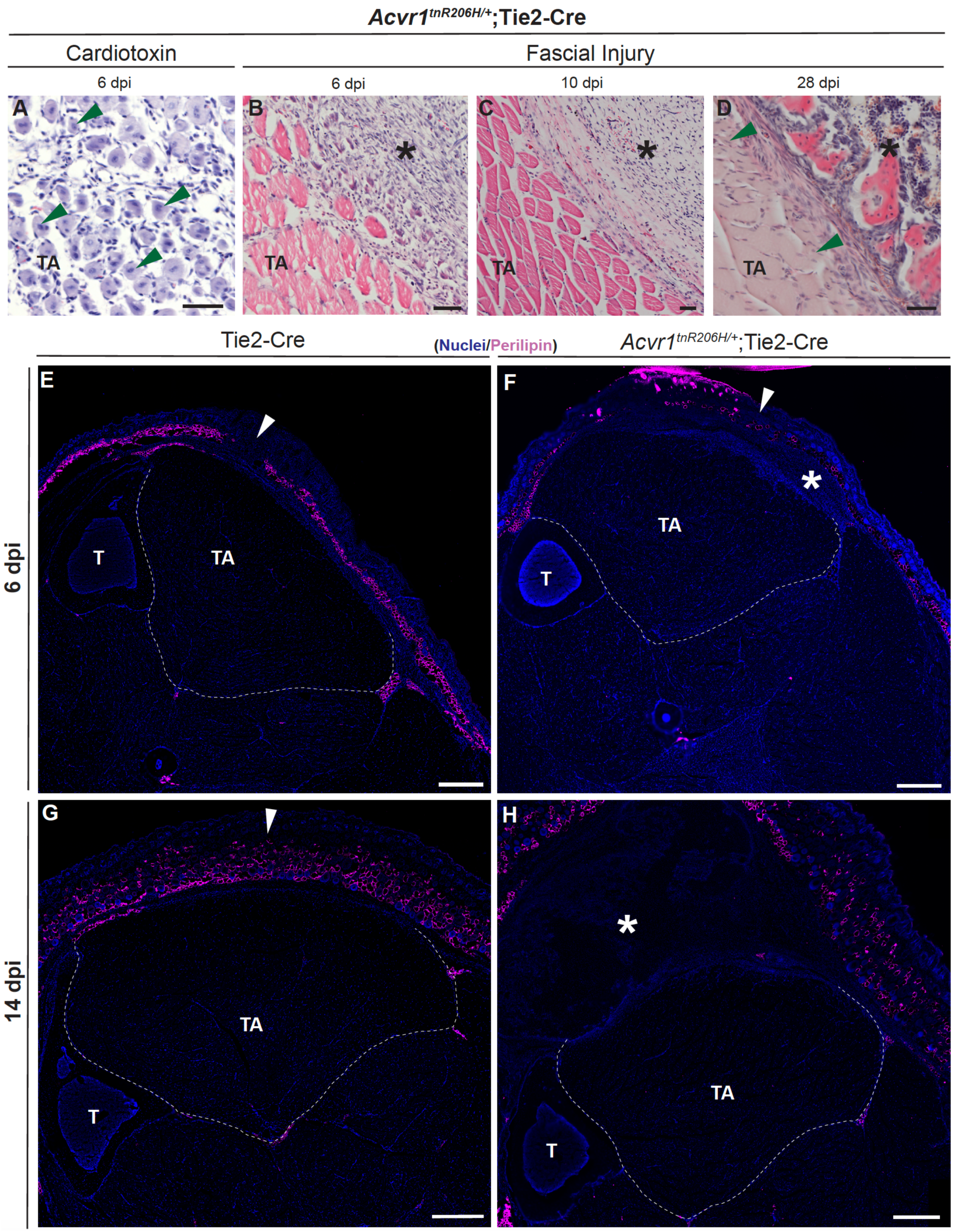
Degenerating muscle adjacent to the site of fascial injury does not regenerate or accumulate adipocytes in *Acvr1^tnR^*^206^*^H/+^*;Tie2-Cre mice. **(A)** H&E-stained cryosections 6 days after cardiotoxin-induced injury of the TA muscle of an FOP mouse (n = 3). Many nascent fibers were present by 6 dpi (identified by central nucleation; several are marked with arrowheads). **(B-D)** H&E-stained cryosections of fascial-injured *Acvr1^tnR^*^206^*^H/+^*;Tie2-Cre mice at 6 (B), 10 (C), and 28 (D) dpi (n = 4, for all time points). Images shown are at the interface of lesional tissue and the remaining, intact, TA muscle. Nascent fibers were absent from hypercellularized regions at 6 and 10 dpi and were only occasionally observed at the muscle/lesion interface at 28 dpi. A small nascent fiber in (D) is identified with an arrowhead. **(E-H)** Fluorescent images of cryosections from control (E, G) and FOP (F, H) mice at 6 (E, F) and 14 (G, H) dpi stained for Perilipin (magenta). Adipocytes were not observed in lesional areas and were only occasionally observed within the remaining intact TA musculature. Cutaneous adipocytes were abundant in both experimental and control mice. The posterior and medial portion of the TA muscle is outlined with a dashed line. Arrowheads mark the approximate site of fascial injury. Asterisks denote areas of hypercellularity (B, C, F) or HO (D, H). T, tibia; TA, tibialis anterior. Female mice were used for all experiments. Scale bars in (A-D) = 50 µm. Scale bars in (E-H) = 500 µm.

*Acvr1^tnR^*^206^*^H/+^*;*Pdgfra^CreERT^*^2^*^/+^* mice displayed a similar muscle degenerative phenotype (Supplemental Fig. 4A, B) with accompanying hypercellularity and abundant FAPs, identified with the Cre-dependent GFP allele, *R26^NG^* ^40^. These data demonstrate that *Acvr1^R^*^206^*^H^*-expressing FAPs are primary drivers of both HO and muscle degeneration in FOP mice. We note that some SOX9+ cells did not express the GFP lineage marker (Supplemental Fig. 4C). These cells could represent FAPs that were recombined at the *Acvr1^tnR^*^206^*^H^* allele but not the *R26^NG^*allele. Alternatively, SOX9+GFP-cells could represent the participation of non-FAP cells in skeletal lesion formation^14^. However, the great majority of these cells were tdTomato-negative (recombined at the *Acvr1^tnR^*^206^*^H^* locus), and therefore likely of FAP origin.

### Muscle degeneration does not stimulate a productive regenerative response in FOP mice

In wild-type mice, pinch- or cardiotoxin-induced muscle injury elicits a robust and rapid regenerative response, with scattered nascent muscle fibers observed by 3 dpi and abundant regenerated fibers by 6 dpi (Fig. 4A)^14,53,56^. In fascial-injured *Acvr1^tnR^*^206^*^H/+^*;Tie2-Cre mice, regenerated fibers were absent from hypercellular regions through 10 dpi. At 28 dpi, nascent fibers were only rarely present, typically at the edge of skeletal lesional tissue (Fig. 4D). Taken together, the data demonstrate that the encroachment of lesional tissue into the TA muscle compartment in FOP mice is associated with pronounced muscle degeneration and lack of a significant muscle regenerative response.

Accumulations of adipocytes and collagen-rich fibrotic tissue are prominent pathophysiological features of most muscular dystrophies and mouse models of impaired regeneration^14,53,56–59^. Whether a fibrotic response was associated with failed regeneration in this model could not be assessed because pre-skeletal and skeletal lesions stain intensely for fibrillar collagen (data not shown). Therefore, this analysis focused on adipogenesis. Notably, no increase in Perilipin+ adipocytes was observed in areas of muscle degeneration at 6 dpi (Fig. 4E-H; Supplemental Fig. 5A), a time point at which adipocyte accumulations are already observed in distinct mouse models of disrupted regeneration^43,60–63^. By 14 dpi, formerly hypercellular regions within the TA muscle compartment were largely contained skeletal lesional tissue (Fig. 1F-H; Fig. 4D). Adipocytes embedded within the remaining TA muscle were rarely observed, and a survey of lesion-spanning tissue sections indicated no increase in adipocytes compared to fascial-injured controls (Fig. 4E-H; Supplemental Fig. 5B)^15^.

### Widespread growth of HO distant from the site of fascial injury in exacerbating models of HO in FOP mice

The highly localized and predictable anatomical position of HO development after the standard fascial injury provided the opportunity to investigate mechanisms of HO growth under conditions in which HO growth is exacerbated. In previous studies, we showed that HO volume is increased dramatically after pinch- or cardiotoxin-induced muscle injury when FOP mice lack the wild-type copy of *Acvr1* (*Acvr1^tnR^*^206^*^H/Flox^*;Tie2-Cre)^14^ or when *Acvr1^tnR^*^206^*^H/+^*;Tie2-Cre mice were treated with JAB0505, an antibody agonist of ACVR1(R206H)^15^. FOP mice that received a small fascial incision and were either treated with a single IP dose of 10 mg/kg JAB0505 at the time of injury or lacked the wild-type *Acvr1* allele exhibited a significant increase in HO volume by μCT at 14 dpi (Fig. 5A-C). Notably, histological analysis of whole lower hindlimb transverse sections using picrosirius red, fast green, and alcian blue (RGB) trichrome staining^64^ suggested that HO lesional tissue largely originated in intermuscular fascia and spread in fascial planes, and directionality of growth could be inferred by the maturity of lesional tissue. Thus, the apparent site of HO initiation close to the fascial injury site was occupied by lesional tissue of maturing bone, whereas apparent leading edges of growing lesions contained cartilage and regions of hypercellularity, which typify lesional areas prior to the appearance of definitive cartilage (Fig. 5D-G). This phenomenon has been previously described as “creeping HO”^65^ and is associated with spontaneous HO^65,66^. These models of exacerbated HO were characterized by foci of HO distant from the injury site, likely representing additional sites of HO initiation within the fascia (Fig. 5D-G). These foci also exhibited gradients of cell maturation such that the directionality of growth could be inferred (Fig. 5D, E). As with untreated heterozygous FOP mice following fascial injury, lesional tissue invaded nearby muscle tissue, which underwent degeneration but failed to undergo a productive regenerative response. In these exacerbated HO models, however, muscle destruction was far more extensive (Fig. 5D-G) and mechanisms that limit the extent of HO growth and muscle destruction following fascial injury of heterozygous FOP mice are apparently inoperative or substantially impaired.

**Figure 5.**
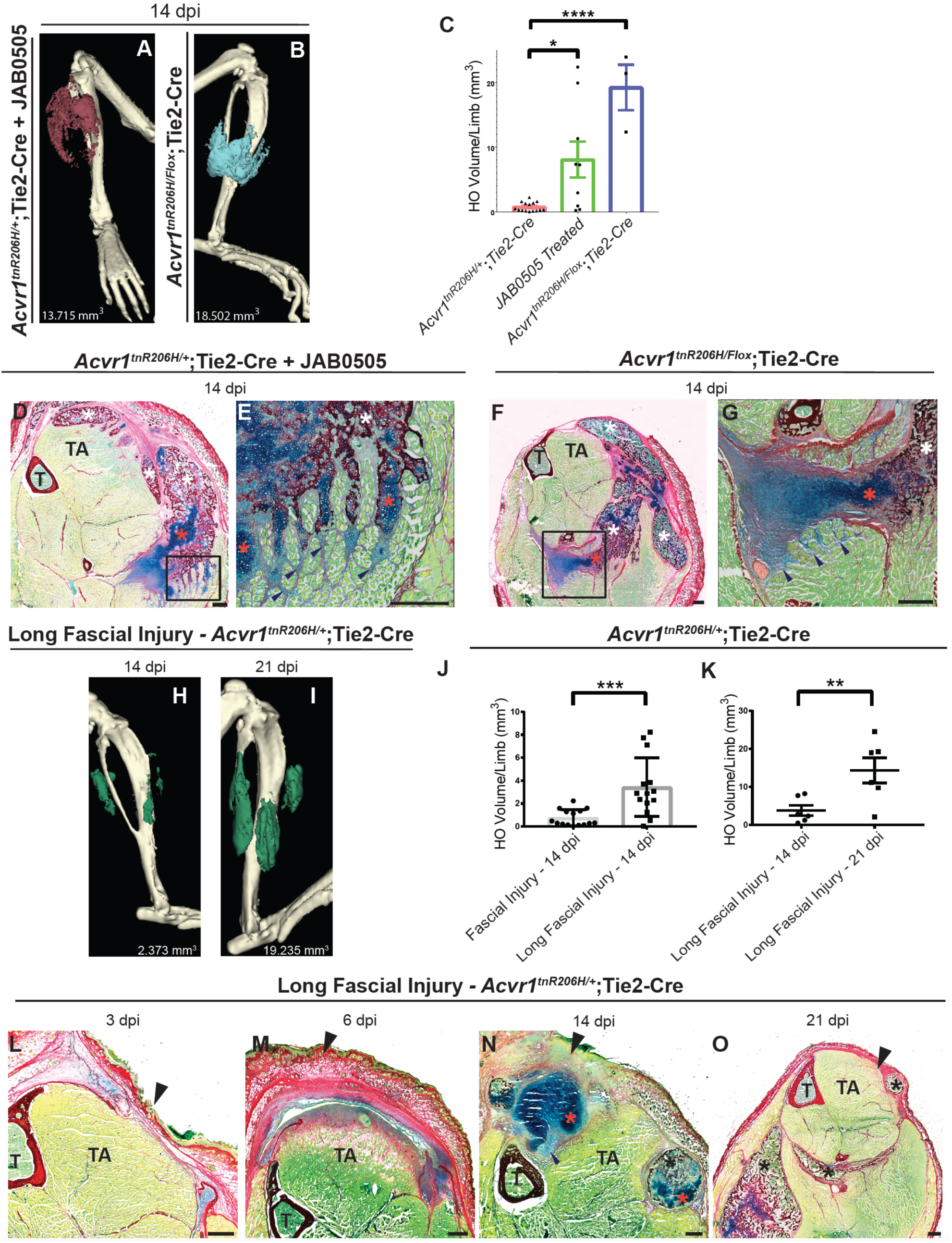
Lesional tissue creeps through the fascia and invades adjacent musculature in exacerbating models of FOP. **(A, B)** µCT images showing exacerbated HO in an *Acvr1^tnR^*^206^*^H/+^*;Tie2-Cre mouse treated with the ACVR1 agonist JAB0505 (A; n = 9), and an FOP mouse lacking a WT *Acvr1* allele (B; *Acvr1^tnR^*^206^*^H/Flox^*;Tie2-Cre; n = 3). HO was pseudocolored maroon or blue. HO responses varied considerably, and images were chosen that showed comparable levels of HO. **(C)** Quantification of HO volumes in *Acvr1^tnR^*^206^*^H/Flox^*;Tie2-Cre mice (n = 3), and *Acvr1^tnR^*^206^*^H/+^*;Tie2-Cre mice with (n = 9) and without (n = 15) JAB0505 treatment. HO volumes were quantified at 14 dpi. **(D-G)** Cryosections of a portion of the lower hindlimb at 14 dpi from either an *Acvr1^tnR^*^206^*^H/+^*;Tie2-Cre mouse treated with JAB0505 on Day 0 (D, E), or an *Acvr1^tnR^*^206^*^H/Flox^*;Tie2-Cre mouse (F, G). Sections were stained with picrosirius red, fast green, and alcian blue (RGB stain). White asterisks show regions of mineralized bone (red). Orange asterisks identify areas of definitive cartilage (dark blue). (E) and (G) correspond to the boxed areas of (D) and (F), respectively. In (E) and (G), finger-like projections of lesional tissue have invaded muscle tissue. Arrowheads point to presumptive leading edges of lesional growth. Areas of hypercellularity stain light blue and skeletal muscle stains green. **(H, I)** Representative µCT images of an *Acvr1^tnR^*^206^*^H/+^*;Tie2-Cre FOP mouse taken 14 (H) or 21 (I) days after a long fascial injury (n = 6). HO is pseudocolored green. (**J)** Quantification of HO volumes induced by a standard (n = 15) or long (n = 14) fascial injury of *Acvr1^tnR^*^206^*^H/+^*;Tie2-Cre mice. µCT images were taken at 14 dpi. **(K)** Longitudinal comparison of HO volumes at 14 and 21 dpi of *Acvr1^tnR^*^206^*^H/+^*;Tie2-Cre mice that received a long fascial injury (n = 6)**. (L-O)** Examples of pre-skeletal and skeletal lesions at 3 through 21 dpi induced by long fascial injury of *Acvr1^tnR^*^206^*^H/+^*;Tie2-Cre mice (n = 4 per time point). RGB-stained cryosections are of the anterior portion of the lower hindlimb. The effects of the large fascial injury largely phenocopied the exacerbated HO phenotypes described above. Black asterisks show regions of mineralized bone. Orange asterisks are cartilaginous areas. Arrowheads mark the approximate location of the fascial injury. Scale bars in (D, F, L-O) = 500 µm. Scale bars in (E, G) = 200 µm. Female mice were used for all experiments. T, tibia; TA, tibialis anterior. Bars in (C, J, K) represent ± SEM. *p ≤ 0.05 **p ≤ 0.01, ***p ≤ 0.001, ****p ≤ 0.0001.

Although the above phenotypes are striking, the models either utilized a genotype that has not been observed in the human population or used pharmacological means of exacerbating responses to injury. Therefore, we tested whether simply increasing the length of the fascial incision in *Acvr1^tnR^*^206^*^H/+^*;Tie2-Cre FOP mice would increase the severity of the response. We produced an incision in the fascia in an identical manner as above, except that the incision encompassed approximately 75% of the TA longitudinal axis (referred to as a long fascial injury). As with the standard fascial incision, control mice showed no indication of muscle damage at day 0, or muscle degeneration at 6 dpi (data not shown). Notably, both HO volume and muscle destruction were dramatically increased following the long fascial incision procedure compared to the standard fascial injury (Fig. 5H-J). HO burden in the lower hindlimb increased at least through 21 dpi (Fig. 5K; Supplemental Fig. 6). This model shares histological features with the other models of exacerbated HO (Fig. 5D-G), including the “creeping” of HO through fascial planes, and initiation of HO at multiple, apparently independent, foci (Fig. 5L-O). Enhancement of HO and the consequent pronounced muscle destruction will be useful for investigating the underlying basis of muscle degeneration and failed regeneration.

## Discussion

Fascia is a network of connective tissue that encloses and supports muscle contraction and stability^67–69^. Fascia plays a critical role in muscle regeneration and wound repair by acting as a scaffolding, a source of extracellular matrix (ECM) remodeling proteins, and by activating FAPs and fibroblasts that remodel and regenerate the ECM^70,71^. The present data indicate that the fascia also serves a primary role in pathological bone formation in FOP. Studies of injury-induced HO in mouse FOP models largely employ direct muscle injury by chemical or physical means to induce HO formation. In normal muscle, these injury modalities cause severe tissue damage and stimulate a robust muscle regenerative response, which includes mobilization of FAPs – multipotent mesenchymal cells dispersed throughout the muscle endomysium that serve positive effector functions in regeneration^72^. Notwithstanding that muscle injury can be an effective means of triggering HO, clear evidence for the initiation of HO intramuscularly is lacking, and previous data^14,17^ suggest an important role for intermuscular fascia in HO initiation and growth. Thus, we previously showed that in a model of spontaneous HO, lesional growth tends to follow fascial planes^14^. In addition, the posterior septum of the lower hindlimb is a common site of HO formation following cardiotoxin injection into the TA muscle^17^. Here, using both Tie2-Cre^28^ and *Pdgfra^CreERT^*^2^ ^29^ drivers to target FAPs, we showed that fascial injury alone, which caused no discernable damage to the TA muscle inferior to the site of injury, is sufficient to induce HO. Together with previous work^15^, the current study supports the conclusion that muscle injury is neither necessary nor sufficient for HO induction.

Evidence for the insufficiency of muscle injury for HO induction derives from studies in which intramuscular injection of methylcellulose caused a localized injury that was sufficient to damage muscle fibers, activate satellite cells, and elicit a muscle regenerative response, but did not induce HO in *Acvr1^tnR^*^206^*^H^*^/+^;Tie2-Cre FOP mice (ref.^15^ and unpublished observations). Thus, although FAPs are interspersed throughout the endomysium, the fine connective tissue layer that surrounds and separates individual muscle fibers, endomysial FAPs appear resistant to osteogenic reprogramming, at least in some contexts. This is unlikely due to differences in developmental potential between FAPs resident in distinct connective tissue layers, as PDGFRα-positive cells from skeletal muscle that does not ossify (e.g., tongue), or even from non-muscle tissues, can form bone when transplanted (unpublished observations). Other parameters that may explain different propensities of muscle-associated tissues to undergo FAP-directed HO include possible differences in FAP density, and distinct signaling and physical environments that may promote or impede HO formation and growth. Understanding the comparative resistance of progenitors within intramuscular tissue to undergo pathological bone formation may suggest approaches to mitigate HO formation in more susceptible tissues.

Hypercellularity of tissues surrounding the fascial injury area was evident by 3 dpi and pronounced by 6 dpi, when a substantial portion of the hyperplastic region included anatomical regions formerly occupied by TA muscle fibers. We showed that FAPs constitute a substantial portion of these early cellular accumulations, and EdU staining (unpublished observations) has shown that many FAPs in these areas of hypercellularity are actively proliferating. FAPs are resident in the injured fascial layers as well as in the endomysium of the underlying TA muscle, and the relative contribution of these FAP populations, whether through proliferation or cell migration, is unclear. Of note, in uninjured muscle tissue, approximately 10% of FAPs carry a recombined *Acvr1^tnR^*^206^*^H^* allele when the Tie2-Cre driver is used^14^, whereas the majority of FAPs in cellular accumulations at 6 dpi were recombined. The proportion of recombined FAPs also increased in *Acvr1^tnR^*^206^*^H/+^*;Tie2-Cre mice following muscle pinch injury^18^. Differential accumulation of R206H-FAPs could result from increased proliferation or decreased apoptosis relative to unrecombined FAPs, or represent continued recombination of *Acvr1^tnR^*^206^*^H^* alleles in previously unrecombined FAPs, perhaps because of increased Tie2-Cre expression after injury. Compared to normal regeneration, FAPs in injured muscle of FOP mice exhibit significantly decreased apoptosis^24^. However, since Cre-dependent *Acvr1^R^*^206^*^H^* expression was not specifically targeted to FAPs in that study, further investigation is required to determine whether decreased apoptosis represented a cell-autonomous effect of *Acvr1^R^*^206^*^H^* in mutant FAPs.

After the standard fascial incision, lesional tissue developed in a predictable anatomical location, directly beneath the incision site. In our experience, the site of HO formation can vary substantially in traditional muscle injury models, complicating studies in which enrichment of progenitors actively engaged in lesion formation is a priority. The predictable location of lesional tissue following fascial injury, and the ability to distinguish unrecombined from recombined cells using the *Acvr1^tnR^*^206^*^H^* allele^14^, will facilitate molecular and biochemical studies of FAP-driven HO. Depending on study objectives, it may also be beneficial to use female FOP mice, as they exhibit greater disease penetrance and HO volumes than males, as shown here and previously^18^.

HO formation within muscle tissue and the consequent effects on muscle structure and function are clinically significant FOP disease manifestations^73,74^. Mechanisms underlying muscle degeneration, however, have been difficult to study in FOP mouse models because the chemical and physical methods used to stimulate HO purposely cause severe muscle damage. The finding that injury to the superficial fascia alone in FOP mice causes severe muscle destruction in the vicinity of the fascial injury now provides an approach for studying the relationship between progenitor-driven HO and muscle degeneration. The catabolic effects of R206H-FAPs on nearby muscle tissue reached a peak at approximately 6 days after fascial injury, a time point at which SOX9+ presumptive cartilage progenitors were abundant in lesional areas, but prior to the appearance of histologically identifiable cartilage. Subsequent time points were characterized by development of definitive cartilage and bone, the growth of which was self-limiting, and stabilization of the remaining muscle tissue juxtaposed to the region of muscle destruction. These observations support the testable hypothesis that loss of muscle integrity is predominantly driven by R206H-FAPs during or shortly after initial stages of R206H-FAP recruitment, when R206H-FAPs become reprogrammed for skeletogenesis. This could explain the extensive and ongoing muscle destruction in exacerbated HO models in which the protracted period of lesional growth is presumptively driven, at least in part, by sustained recruitment of R206H-FAPs^15^ as growing lesional tissue “creeps” through the intermuscular fascia. Gene expression studies at stages of R206H-FAP reprogramming may identify direct or indirect effectors of muscle destruction.

Previous studies have shown that muscle regeneration is impaired in FOP mice following muscle injury when either a globally expressed Cre driver^24^ or Tie2-Cre^14^ was used. Further, expression of *Acvr1^R^*^206^*^H^*in cultured satellite cells significantly reduced their differentiation capacity^24^. These data are consistent with the notion that regeneration deficits result from both cell-autonomous and cell-non-autonomous effects of *Acvr1^R^*^206^*^H^*expression in satellite cells and FAPs, respectively. Interestingly, muscle regeneration was reportedly not impaired when *Acvr1^R^*^206^*^H^* expression in mice was exclusively targeted to satellite cells^24^, indicating that the genetic status of FAPs is the primary determinant of regenerative outcomes, a conclusion supported by FAP-satellite cell co-culturing experiments^24^. The current study strongly implicates R206H-FAPs not only in muscle destruction, but also in the abrogation of muscle’s regenerative capacity, as no nascent fibers were observed in hypercellular regions formerly occupied by TA muscle fibers. This represented an apparent loss of regenerative capacity rather than delayed regeneration, as regenerated muscle fibers were absent at all stages examined, including 14 dpi, when regeneration is essentially complete after muscle injury of wild-type mice. It is notable that in co-culturing experiments, R206H-FAPs had only a modest (but statistically significant) inhibitory effect on differentiation of wild-type satellite cells^24^. This is in contradistinction to the far more pronounced effects in vivo reported here, in which FAPs, but not satellite cells, were targeted for *Acvr1^R^*^206^*^H^* expression. These data suggest that the inhibitory effects of R206H-FAPs on satellite cells may be largely indirect. In normal regeneration, FAPs function as positive effectors of regeneration at multiple levels that include direct interactions with satellite cells to promote their differentiation, maintenance of the satellite cell pool, conditioning of the ECM, secretion of cytokines, and other functions^72,75^. Transcriptomic analyses of FAPs and satellite cells in FOP mice should identify candidate genes and pathways responsible for impaired muscle regeneration in FOP mice.

Most models of impaired regeneration, regardless of the proximate cause, are characterized by accumulations of adipocytes and fibrotic tissue, which can interfere with muscle structure and function^14,53,56–59^. Available evidence indicates that FAPs are the predominant source of these non-muscle accumulations^72,75^. An important aspect of the non-regeneration phenotype reported here is the absence of adipocyte accumulations (fibrosis could not be evaluated) in areas of muscle degeneration. R206H-FAPs were abundant in hypercellular regions, and these *Acvr1^R^*^206^*^H^*-expressing FAPs were likely destined for skeletogenic commitment and differentiation. However, when either the Tie2-Cre or *Pdgfra^CreERT^*^2^ driver was used, unrecombined FAPs (and therefore not expressing the *Acvr1^R^*^206^*^H^*allele) were easily identifiable in hypercellular regions at the approximate stage at which muscle degeneration was maximal (∼ 6 dpi). Unrecombined FAPs do not appreciably contribute to HO lesions induced by direct muscle injury^14^ or fascial incision (data not shown) and would be expected to retain an intrinsic capacity for adipogenic differentiation. A better understanding of FAP biology in the fascial injury model may suggest strategies for preventing adipocyte accumulations in other diseases and conditions.

## Acknowledgements

We thank members of the Goldhamer Lab for critical feedback during the course of this work, Brenden Griffith, Heather Jamieson, and Ingrid Schwarz for careful review of the manuscript, and Alexion Pharmaceuticals for providing JAB0505. This work was supported by a grant from the NIH (R01AR072052) and University of Connecticut institutional funds.

