## Supplemental Figures for "The role of muscle fascia in heterotopic ossification and maintenance of skeletal muscle integrity in fibrodysplasia ossificans progressiva"

### Supplemental Figures and Legends

**Supplemental Figure 1. Depiction of fascial surgery method and incidence of HO in experimental and sham-injured mice. (A)** Example of a typical skin incision made superficial to the TA muscle to access the underlying superficial fascia and epimysium. **(B)** Image of the standard fascial injury, which consisted of a 2-3 mm cut to the fascia along the longitudinal access of the TA muscle at the approximate mid-belly. In the image shown, the skin above the TA muscle was removed to more easily visualize the fascial injury. Arrowheads denote the boundaries of the cut in the fascia. **(C)** Comparison of the frequency of HO formation following sham (skin incision only) or fascial injury of *Acvr1*$^{tnR206H/+}$;Tie2-Cre mice. Mice were scored by µCT for HO at 14 dpi (n = 11 for both sham and experimental groups). Both male and female mice were used. **(D)** Representative example of HO formation in the subset of *Acvr1*$^{tnR206H/+}$;Tie2-Cre mice that formed HO after sham injury. The image is an H&E-stained transverse cryosection of the entire lower hindlimb 14 days after sham injury. Arrows denote the undisturbed fascia that separates the HO (asterisk) from the underlying TA muscle. The skin was removed prior to fixation. T, tibia; TA, tibialis anterior. Scale bar = 500 µm.

**Supplemental Figure 2. HO formation after fascial injury of FOP mice is highly penetrant but HO volumes show considerable variation. (A)** Comparison of HO volumes of male (n = 8) and female (n = 15) *Acvr1*$^{tnR206H/+}$;Tie2-Cre mice and female *Acvr1*$^{tnR206H/+}$*;Pdgfra*$^{CreERT2/+}$ mice (n = 5) at 14 dpi. **(B)** Longitudinal analysis of HO formation in female *Acvr1*$^{tnR206H/+}$;Tie2-Cre mice through 4 weeks after fascial injury. Colored lines indicate individual mice (n = 4) and data points represent the average HO volume of both limbs. **(C-D)** µCT images of HO in a female *Acvr1*$^{tnR206H/+}$;Tie2-Cre mouse at 21 (C) and 28 (D) dpi. Images shown are derived from the mouse represented by the green line in (B).

**Supplemental Figure 3. Muscle fiber counts in control and FOP mice. (A, B)** Quantification of myofibers in the TA and EDL muscles of uninjured *Acvr1*$^{tnR206H/+}$;Tie2-Cre mice, fascial-injured *Acvr1*$^{tnR206H/+}$;Tie2-Cre mice, and fascial-injured Tie2-Cre (control) mice at 14 (A) and 6 (B) dpi. Sections were chosen at anatomical positions that represented the position of maximal lesional size, and fibers were quantified on one section from each of four mice for each group. Muscle fiber counts of uninjured FOP mice and injured control mice of the same table row were at matching anatomical positions to control for position-dependent differences in fiber numbers along the longitudinal axis of the lower hindlimb.

**Supplemental Figure 4**. **Lesion formation and muscle degeneration following fascial injury in *Acvr1*$^{tnR206H/+}$;*Pdgfra*$^{CreERT2/+}$;*R26*$^{NG/NG}$ FOP mice**. The injury response largely phenocopies that of FOP mice in which the Tie2-Cre driver is used. **(A)** H&E-stained cryosection of the anterior portion of the lower hindlimb. Asterisks mark lesional areas. **(B, C)** Fluorescent images of a cryosection from an *Acvr1*$^{tnR206H/+}$;*Pdgfra*$^{CreERT2/+}$;*R26*$^{NG/NG}$ mouse at 6 dpi. The panels in (C) correspond to the boxed area in (B). In some lesional areas, the great majority of GFP+ cells are also SOX9+ (three of many examples shown at arrows in C). The left image in (C) is a merged image showing colocalization of GFP and SOX9. The middle and right panels are single-channel images. The arrowhead in (A) marks the approximate site of fascial injury. T, tibia; TA, tibialis anterior. Female mice were used for all experiments. Scale bars in (A), (B), and (C) are 500 µm, 200 µm, and 50 µm, respectively.

**Supplemental Figure 5. Degenerating muscle close to the site of fascial injury does not accumulate adipocytes in *Acvr1*$^{tnR206H/+}$;*Pdgfra*$^{CreERT2/+}$;*R26*$^{NG/NG}$ mice. (A, B)** Fluorescent images of cryosections from FOP mice at 6 (A) and 14 (B) dpi stained for Perilipin (magenta). Adipocytes were not observed in lesional areas and were only occasionally observed within the remaining intact TA musculature. Cutaneous adipocytes were abundant in experimental and control (not shown) mice. The posterior and medial portion of the TA is outlined with a dashed line. Green fluorescence driven by the *R26$^{NG}$* allele is not shown to allow better visualization of Perilipin staining. Arrowheads mark the approximate site of fascial injury. Asterisks denote areas of hypercellularity (A), which is characteristic of pre-skeletal lesional tissue, or mark HO (B). T, tibia; TA, tibialis anterior. Female mice were used for all experiments. Scale bars in (A, B) = 500 µm.

**Supplemental Figure 6. Time course of lesional growth following long fascial injury of *Acvr1$^{tnR206H/+}$*;Tie2-Cre and *Acvr1$^{tnR206H/+}$*;*Pdgfra$^{CreERT2/+}$* mice.** In both models, HO volumes continued to increase after 14 dpi. **(A)** Longitudinal study of HO growth through 28 dpi in female *Acvr1$^{tnR206H/+}$*;*Pdgfra$^{CreERT2/+}$* FOP mice (n = 2). **(B)** Longitudinal analysis of HO formation in individual female *Acvr1$^{tnR206H/+}$*;Tie2-Cre mice through 21 dpi. Colored lines indicate individual mice (n = 6).

**Supplemental Figure 1**

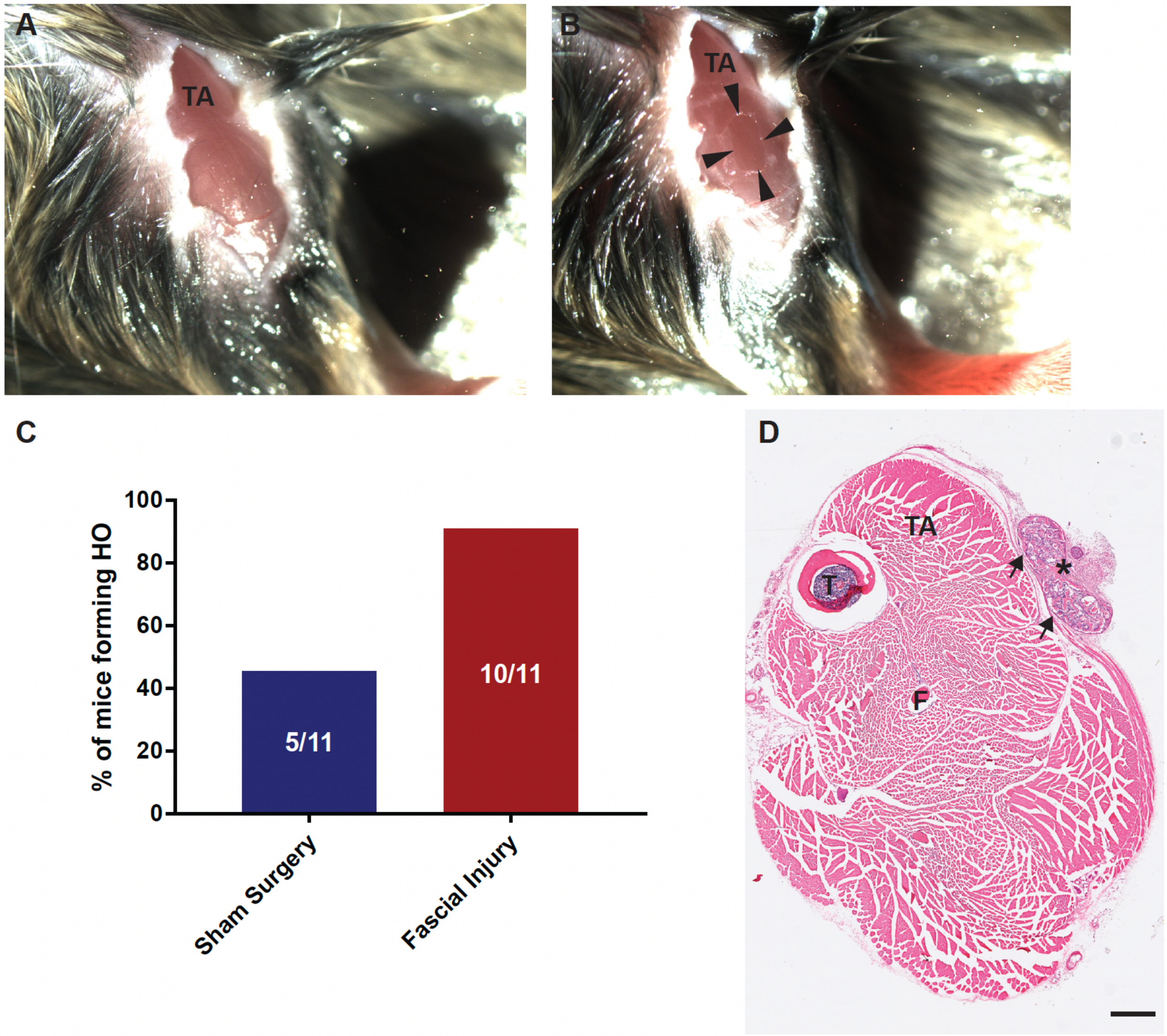

### Supplemental Figure 2

**A**

| HO Volume/Limb (mm$^3$) | | |
|---|---|---|
| *Acvr1* $^{tnR206H/+}$;Tie2-Cre | | *Acvr1* $^{tnR206H/+}$;*Pdgfra* $^{CreERT2/+}$ |
| **Male** | **Female** | **Female** |
| 0.07 | 0.15 | 0.69 |
| 0.17 | 0.04 | 0.11 |
| 0.15 | 1.28 | 0.15 |
| 0.06 | 2.23 | 0.07 |
| 0.00 | 1.59 | 0.29 |
| 0.02 | 1.35 | |
| 0.08 | 1.58 | |
| 0.08 | 0.14 | |
| | 0.10 | |
| | 0.27 | |
| | 0.30 | |
| | 0.47 | |
| | 1.13 | |
| | 0.20 | |
| | 0.29 | |

**B**

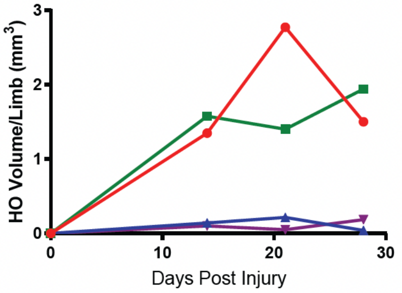


***Acvr1*$^{tnR206H/+}$;Tie2-Cre**

21 dpi

**C**

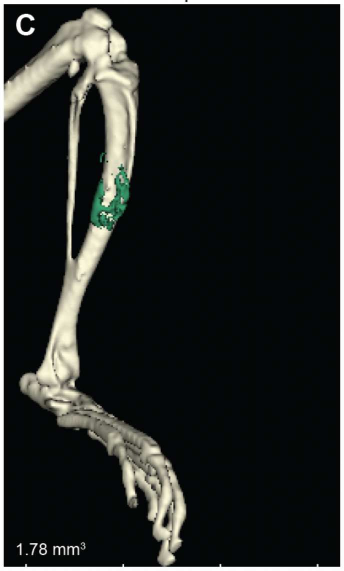


28 dpi

**D**

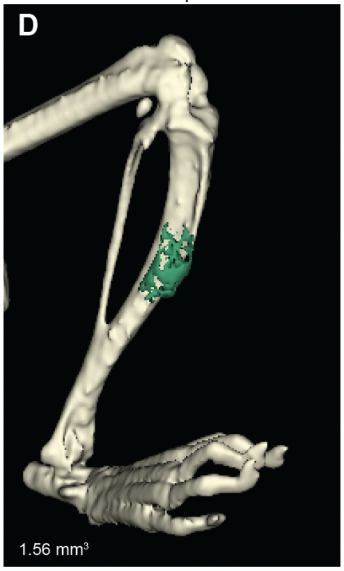

**Supplemental Figure 3**

**A**

### 14 dpi

| Fiber Count in TA and EDL (Percent fibers compared to uninjured) | | |
|---|---|---|
| *Acvr1*$^{tnR206H/+}$;Tie2-Cre | Tie2-Cre | *Acvr1*$^{tnR206H/+}$;Tie2-Cre |
| Uninjured | Fascial Injured | |
| 3640 (100) | 3572. (98.1) | 2019 (55.5) |
| 4071 (100) | 3724 (91.5) | 2798 (68.7) |
| 3902. (100) | 3714 (95.2) | 3666 (93.9) |
| 4133 (100) | 3716 (89.9) | 2126 (51.4) |

**B**

### 6 dpi

| Fiber Count in TA and EDL (Percent fibers compared to uninjured) | | |
|---|---|---|
| *Acvr1*$^{tnR206H/+}$;Tie2-Cre | Tie2-Cre | *Acvr1*$^{tnR206H/+}$;Tie2-Cre |
| Uninjured | Fascial Injured | |
| 2618 (100) | 2741 (104.7) | 1268 (48.4) |
| 3737 (100) | 3649 (97.6) | 2880 (77.1) |
| 3640 (100) | 3685 (101.2) | 2296 (63.1) |
| 3911 (100) | 3944 (100.8) | 3382 (86.5) |

**Supplemental Figure 4**

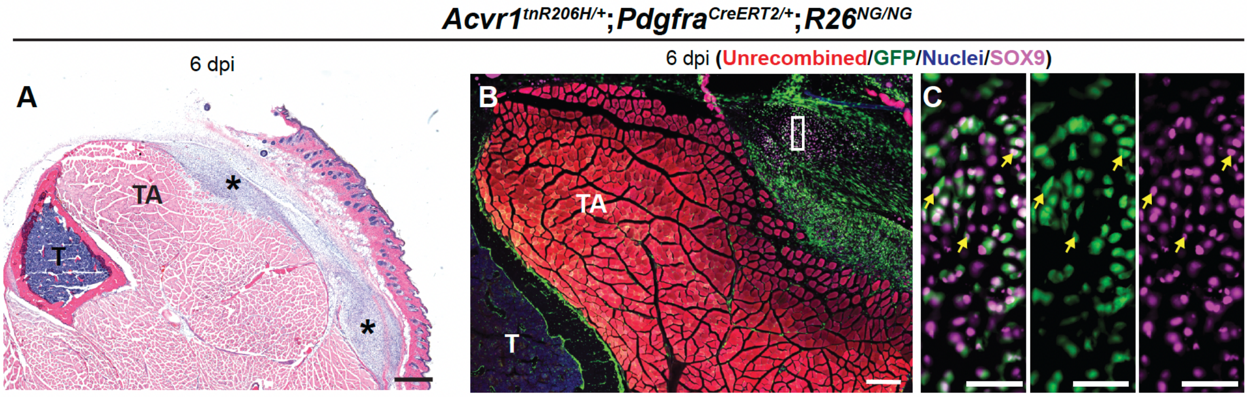

**Supplemental Figure 5**

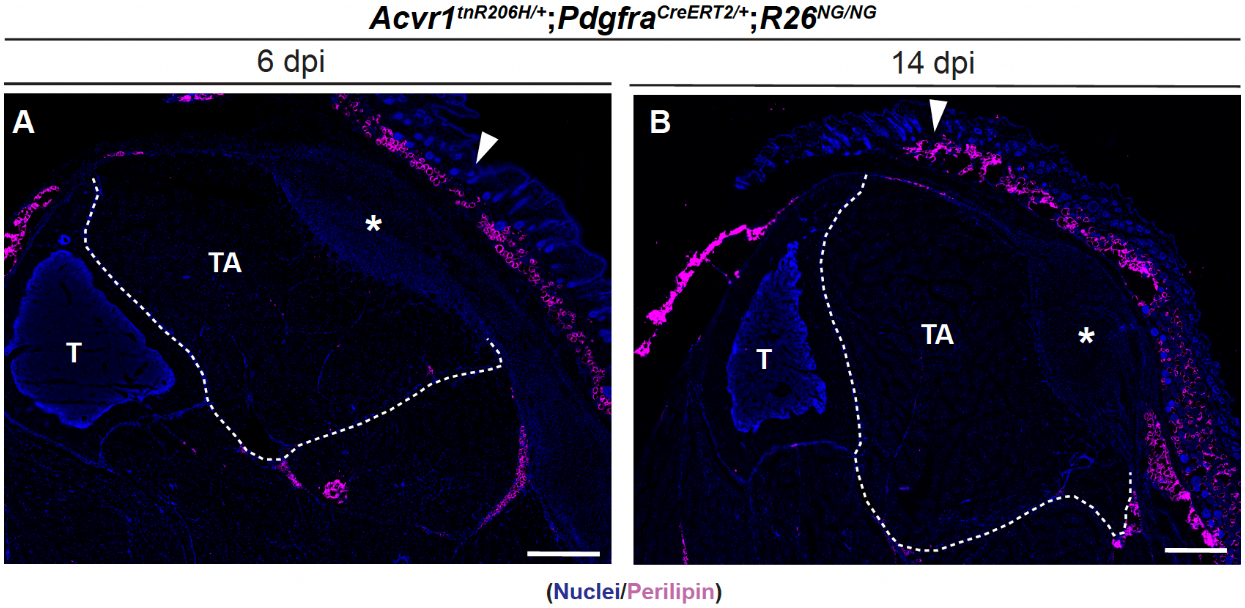

**Supplemental Figure 6**

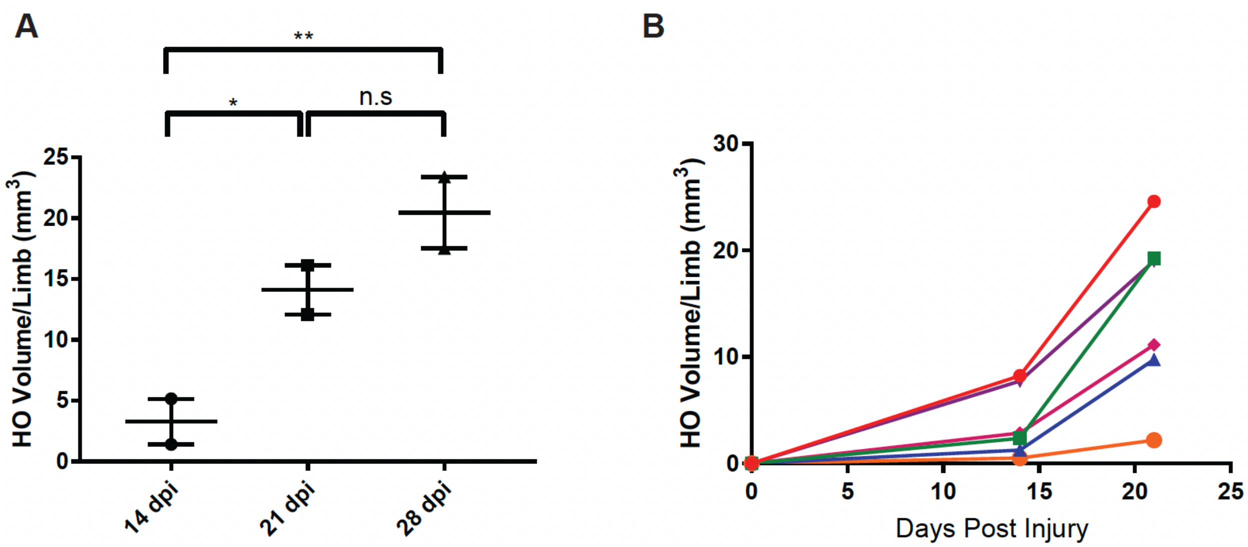
